# Population structure of the *Brachypodium* species complex and genome wide association of agronomic traits in response to climate

**DOI:** 10.1101/246074

**Authors:** Pip Wilson, Jared Streich, Kevin Murray, Steve Eichten, Riyan Cheng, Niccy Aitkin, Kurt Spokas, Norman Warthmann, Accession Contributors, Justin Borevitz

## Abstract

The development of model systems requires a detailed assessment of standing genetic variation across natural populations. The *Brachypodium* species complex has been promoted as a plant model for grass genomics with translational to small grain and biomass crops. To capture the genetic diversity within this species complex, thousands of *Brachypodium* accessions from around the globe were collected and sequenced using genotyping by sequencing (GBS). Overall, 1,897 samples were classified into two diploid or allopolyploid species and then further grouped into distinct inbred genotypes. A core set of diverse *B. distachyon* diploid lines were selected for whole genome sequencing and high resolution phenotyping. Genome-wide association studies across simulated seasonal environments was used to identify candidate genes and pathways tied to key life history and agronomic traits under current and future climatic conditions. A total of 8, 22 and 47 QTLs were identified for flowering time, early vigour and energy traits, respectively. Overall, the results highlight the genomic structure of the *Brachypodium* species complex and allow powerful complex trait dissection within this new grass model species.

## Introduction

Climate change is impacting the production of food worldwide (1) while increasing global demand will soon outstrip the rate of improvement in crop yield by traditional breeding methods (2). To address food and climate security, there is a need for agricultural innovation across a range of scientific disciplines, from genomics to phenomics in new species across the landscape (3). Key to breeding for more variable future climates, as well as for broad adaptability, is understanding the plasticity of the genetic architecture of agronomic traits across environments. The use of controlled growth cabinets, that can mimic regional diurnal and seasonal climate types (e.g. 4), allows us to examine the genetic architecture underlying complex adaptive traits across field like environments.

Two complex traits that have a large impact on yield are ear emergence and early vigour. The timing of ear emergence is crucially important to yield in many grain growing regions, including Australia where early flowering may lead to cold-induced sterility while late flowering may result in heat stress or lack of water limiting grain filling. Early vigour, defined as an increase in the above ground biomass prior to stem elongation, is a beneficial trait in many environment types, especially when combined with increased transpiration efficiency (5). Since vapour pressure is low in winter, increased biomass during early growth improves plant water use efficiency. Early vigour also increases competition against weeds, reduces soil evaporation and may improve yields by increasing total seasonal biomass (6). Energy use efficiency is a relatively understudied component of plant growth that represents the efficient transfer of energy, acquired through photosynthesis, to the grain, and may significantly affect yield. Early studies indicate that energy efficiency, via lower respiration rates, are correlated with an increase in biomass in monocot species (7,8). Identification of the genetic architecture of energy use efficiency, timing of heading and early vigour traits, as well as the genetic sensitivity to future temperature profiles, could accelerate breeding in crop species via selection for improved predicted yields in the future.

Genome Wide Association Studies (GWAS) combine dense genetic markers identified via next generation sequencing and high throughput phenotyping to identify the causative alleles and to predict complex quantitative traits (9). The improvement of crop yield involves many complex traits and the expression of these traits can be highly dependent on the growth environment. GWAS is an excellent method for mapping and predicting yield-related traits and their interaction with the environment. GWAS has been undertaken in a number of crop species; for dozens of agronomic traits in diploid species such as rice, barley and corn (for review see 10), and has even be used reasonably successfully in wheat despite the added complexity of a hexaploid genome (e.g. 11).

*Brachypodium distachyon* is a model species for temperate C3 grass crops such as wheat, barley, rye and oats as it is also located in the Pooideae family and has a number of advantageous characteristics as a model species (12–15). *B. distachyon* also has a number of advantages over the related domestic Pooideae for a GWAS approach as it is a wild species with a wide climatic distribution, resulting in diverse phenotypes, as well as wide genomic diversity, for traits involved in life strategy and abiotic stress tolerance. *B. distachyon* has a fully sequenced small genome of 270Mb (16) compared to the 16Gb of wheat (17) or 5.1Gb of barley (18). It also contains a low percentage of repetitive non-coding DNA at 21.4% of nucleotides compared to more than 80% in wheat (19) and 84% in barley (18). This means that sequence reads from *B. distachyon* are much easier to identify and align compared to wheat, with a larger proportion of the sequencing providing useful reads. Finally, and perhaps most importantly, is the short stature of *B. distachyon* which allows large numbers of plants to be taken through full life cycles in controlled growth conditions.

*Brachypodium* is widespread throughout temperate regions, including its native Mediterranean range and introduced range in Australia, South Africa and western USA (20,21). A large number of accessions have been collected throughout the world by the *Brachypodium* community but the use of these collections in genomic association studies has been delayed by the cryptic nature of the *Brachypodium* species complex. The three species in this complex are difficult to distinguish in the field and include the diploid *B. distachyon*; the diploid *B. stacei*; and the allotetraploid *B. hybridum* which contains one *B. distachyon* -like genome and one *B. stacei* -like genome (22–24). To add to the complexity there is evidence of distinct subgroups or subspecies of *B. distachyon*, (21,25). While the genome of the Bd21 genotype of *B. distachyon* was published in 2010, the genome of *B. stacei* and other SNP corrected genomes were released online in 2016 (DOE-JGI, http://phytozome.jgi.doe.gov/). Recently, a *B. distachyon* pan genome was published identifying geographic diversity and many new genes not identified in the initial reference (26). Prior to our study, species identification has commonly been undertaken by morphoanatomical classification, a small number of markers or cytology (e.g. 17,23). There is a need for a rapid identification of species, subspecies and genotype lineages, within the *Brachypodium* species complex to aid the selection of HapMap sets and to enable landscape genomic studies of migration and adaptation.

In this study, we aimed to 1) characterise the species, genotype and population structure of a *Brachypodium* global diversity set to select a core haplotype mapping set for GWAS in *B. distachyon* and 2) identify the genetic architecture and plasticity of the agriculturally relevant traits of heading date, early vigour and energy use efficiency in response to climate.

## Results

### Cryptic Brachypodium Species, diverse genotypes and population structure identified using Genotyping by Sequencing

To establish a diverse set of germplasm, thousands of *Brachypodium* accessions were collected on trips to south-west Europe, south-eastern Australia, western USA and through collaborations with the international *Brachypodium* community (https://github.com/borevitzlab/brachy-genotyping/blob/master/metadata/brachy-metadata.csv). Out of these, 1968 accessions were grown to produce single seed descent lines in the greenhouses at ANU for subsequent genomic analysis. A reduced representation approach, *PstI* digest, genotyping-by-sequencing (GBS; 26–28) was used to genetically-profile the accessions.

Although once described as a single species, *B. distachyon* has more recently been shown to exist as a species complex consisting of a 5 Chromosome *B. distachyon*, 10 Chromosome *B. stacei*, and a 15 chromosome allopolyploid *B. hybridum* (22). To categorise each accession into species within the *Brachypodium* complex, GBS tags were mapped to a merged reference genome consisting of *B. distachyon* (Bd21-3) and *B. stacei* (ABR114) [v1.1 DOE-JG, https://phytozome.jgi.doe.gov]. Most accessions were readily distinguished as having reads that aligned to either or both reference genomes (see methods; Supp. S01). The majority of accessions, 56% (1100/1968) were identified as *B. hybridum.* In contrast, only 3% (60/1968) were classified as *B. stacei*, while 35% (698/1968) were *B. distachyon*. The remaining 6% (110/1968) could not be definitively assigned. Mapping of the accessions’ geographic locations showed that *B. hybridum* has expanded across the globe representing essentially all the collections outside the native range (Fig 1A). Conversely, *B. distachyon* is largely limited to the native Mediterranean and Western Asian regions, with *B. stacei* in the same area but was less common.

**Figure 1.**
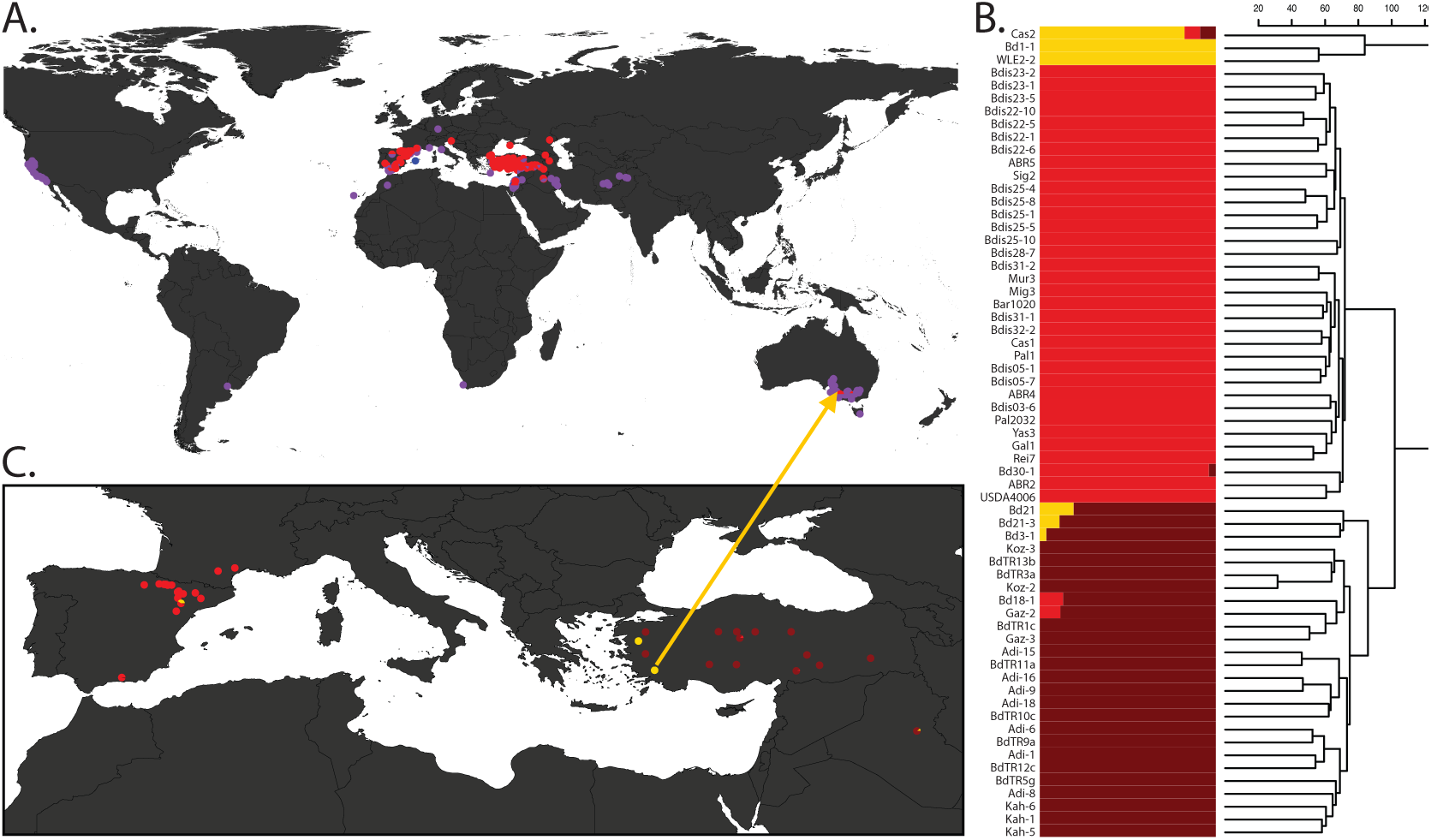
Distribution and Genomic Diversity of the *B. distachyon* complex. A) Geographic distribution of 1573 *Brachypodium* complex accessions classified by species: pink = *B. distacyhon,* blue = *B.stacei* and purple = *B. hybridum* B) Structure plot of the *B. distachyon* species, K=3; and C) Geographic structure of *B. distachyon* across Iberian Peninsula and Turkish region. Proportions of pies represent the number of each *B. distachyon* subgroup (from B) at each site. The arrow from C to A shows the Australia *B. distachyon* (WLE2-2) and the near identical accession from Turkey (BdTR9f).

Due to the highly selfing nature of all *Brachypodium* species, we next sought to categorise accessions into unique whole genome genotypes representing a single inbred lineage. We used the SNPRelate package (31) to cluster 72 genotypes from 490 high quality *B. distachyon* accessions genotyped at 81,400 SNPs (see methods; https://github.com/borevitzlab/brachy-genotyping-notes; Supp. S02). Recombinant inbred lines, included as positive controls, were often called unique genotypes as expected, but were excluded from subsequent analysis of natural population structure.

### Whole genome variation

Whole genome sequencing was performed on a set of 107 *B. distachyon* accessions to determine high density variation at multiple levels, patterns of linkage disequilibrium, and to enable genome wide association studies (GWAS). The samples ranged across the subspecies, population and family levels revealing 2.65M polymorphic sites (1% diversity, θ, on 260Mb). Most samples were the A subspecies with 3 clear outliers (Bd1-1, Cas2, WLE2-2) belonging to the B subspecies. The subspecies showed fixed divergence at 6.5% of sites (169k with <100 expected by chance among 3 accession outliers). Within the A subspecies there were 2 clear populations with 1.5% fixed divergence among groups. Accessions within the same genotypic lineage diverged at between 0.1-to 0.4% of SNPs highlighting the relative amount of rare variation within a unique genotype. A balanced set of representative accessions across the genotype lineages within just the A subspecies was selected for further genomic and phenomic analysis (Supp. S03).

Previous genetic analysis on smaller data sets had shown *B. distachyon* to have substantial levels of population structure (20,25,26,32,33). We sought to refine the ancestral population structure of *B. distachyon* by reducing 107 accessions to 63 highly diverse genotypes using 2,648,921 SNPs. To reduce data complexity SNPs were subsampled to every 100th site to create a final SNP matrix of 26,490 variants that were fed into STRUCTURE v.2.3.4 (34; Supp. S04). STRUCTURE analysis identified three main subgroups among *B. distachyon* genotypes and seven admixed lines (Fig 1B). The yellow lineage was the most diverged and represents subspecies B, with the brown and red structure groups representing the predominantly East and West populations of the A subspecies. To visualise the geographic distribution, the ancestral group composition was summed across accessions for each geographic site (Fig 1C). The single *B. distachyon* accession from Australia WLE2-2 was nearly identical to BdTR9f (GBS data, Supp. S02) from southern western Turkey, where it may have originated from. It is shown in its ancestral location (Fig 1C arrow).

Linkage disequilibrium (LD) was calculated for consecutive windows of 2000 SNPs across the genome. There was large variation in LD across the genome (Supp. S05) with the median LD 113kb (50 – 235kb interquartile range) and the maximum was greater than 2.4Mb.

### Determining the best traits and climatic conditions for GWAS in *B. distachyon*

For our GWAS study we wanted to identify high throughout non-destructive phenotypic measures with high heritability. We also wanted to determine the best environmental conditions to characterise our trait of interest. Hence two preliminary experiments were undertaken, one for flowering time and one for early vigour.

Flowering time was chosen as an ideal trait for GWAS as it has high heritability in many species including Arabidopsis (35)and barley (36). In previous studies of *B. distachyon* it has been found that the dependence of flowering time on vernalisation and photoperiod varies between accessions (37–40). This study aimed to identify QTL for earliness *per se* in flowering, i.e. those responsive to the accumulation of thermal time. Hence a preliminary experiment was undertaken to determine if our conditions could meet the vernalisation requirements of all *B. distachyon* accessions and to determine which accessions had strong vernalisation requirements in our conditions. To do this 266 diverse A and B - subspecies accessions, with 5 accessions replicated 5-6 times were grown in both a simulated winter sowing, starting 01 June, and a spring sowing, starting 01 September, in Wagga Wagga, NSW, Australia (Supp. S06). Ear emergence was monitored, as a surrogate measure for flowering time as flowering occurs largely within the ear in *B. distachyon* so is hard to accurately record (Supp. S07). Out of the 266 accessions there were 17 accessions that did not flower in the Spring condition, indicating a strong vernalisation requirement (Supp. S08A). All lines flowered in the Winter condition, indicating that night temperatures of 4°C were sufficient to meet vernalisation requirement. As expected, days to ear emergence showed a strong heritability in the winter condition, as calculated from the replicated lines (*h*^2^=0.96). The thermal time to flowering was calculated to determine the dependence of flowering on the accumulation of thermal time. The fast cycling accessions, that were not vernalisation requiring, still required a larger thermal time accumulation than the vernalisation requiring accessions (Supp. S08B). This indicates that these either have some low-level requirement for vernalisation that is not being fully met in the Spring condition or that the photoperiod is also a factor in this relationship. As this study aimed to identify QTL for earliness *per se* in flowering, i.e. those responsive to the accumulation of thermal time, we attempted to exclude vernalisation and photoperiod effects by focusing on the winter condition for the GWAS experiment.

In temperate grass crops such as wheat and barley, early vigour can result in an increased yield in short seasons or in seasons where there is high rainfall (reviewed in 6). Often the dimensions of seedling leaves are measured as a nondestructive surrogate measure for early vigour (41,42). To confirm that this was also an appropriate surrogate measure for early vigour in *B. distachyon,* a highly replicated (n = 10) validation experiment was performed on six diverse *B. distachyon* lines (Supp. S09A) in a simulated Wagga Wagga, seasonal climate starting on 01 September (Spring). After seven weeks, when plants had between four and five mainstem leaves, the dimensions of leaf #3, seedling height, total leaf area, and above ground dry weight were measured and phenotypic correlations were calculated (Supp. S09B). Narrow sense heritability was also calculated to determine which early vigour trait would provide the most power for mapping QTLs with GWAS (Supp. S09C). Leaf #3 width and length correlated well with above ground biomass (r^2^=0.46, p<0.01 and r^2^=0.48, p<0.01, respectively) and had quite high heritabilities of *h*^2^=0.60 and *h*^2^=0.64, respectively, as compared to above ground dry mass, *h*^2^=0.51. Interestingly, seedling height also had a strong correlation with above ground biomass (r2=0.74, p<0.01) with a heritability of *h*^2^=0.74. However, this trait was also more highly correlated with development, as indicated by the number of leaves (r2=0.21, p<0.01), than the dimensions of leaf #3. To get the most direct measure of early vigour the dimensions of leaf #3 were chosen as the focus for the GWAS.

### Selection of global HapMap set

High-level population structure confounds GWAS when there are few segregating SNPs in common between ancestral groups relative to variation within each subgroups (43). Here we focused on subspecies A which contains a majority of unique genotypes, resulting in a HapMap set of 74 genotypes. Within the A subspecies there is still clear population structure, but further subset selection would limit both the sample size and the phenotypic and genotypic diversity, reducing the rate of true positive results. This residual relatedness between lines was accounted for by including a kinship matrix in the GWAS model.

### Early vigour and ear emergence shows genotypic variation in response to different simulated environments

To determine the genetic architecture for ear emergence date, early vigour, and a range of other agronomic traits, the refined and balanced HapMap set of 74 *B. distachyon* accessions (Supp. S10), with four biological replicates, were grown in two simulated conditions in specially modified growth chambers (4). To determine the effect of an increase in temperature in line with climate change predictions on the traits of interest, the conditions modelled a present (2015, Fig 2A) and a future (2050, Fig 2B) temperature profile at Wagga Wagga, NSW, Australia. The appropriate increase in average maximum and minimum temperature for each month was determined using an average of 12 global climate change models determined to be high confidence for south east Australia using the Climate Futures Tool (Fig 2C, 42; Supp. S11).

**Figure 2.**
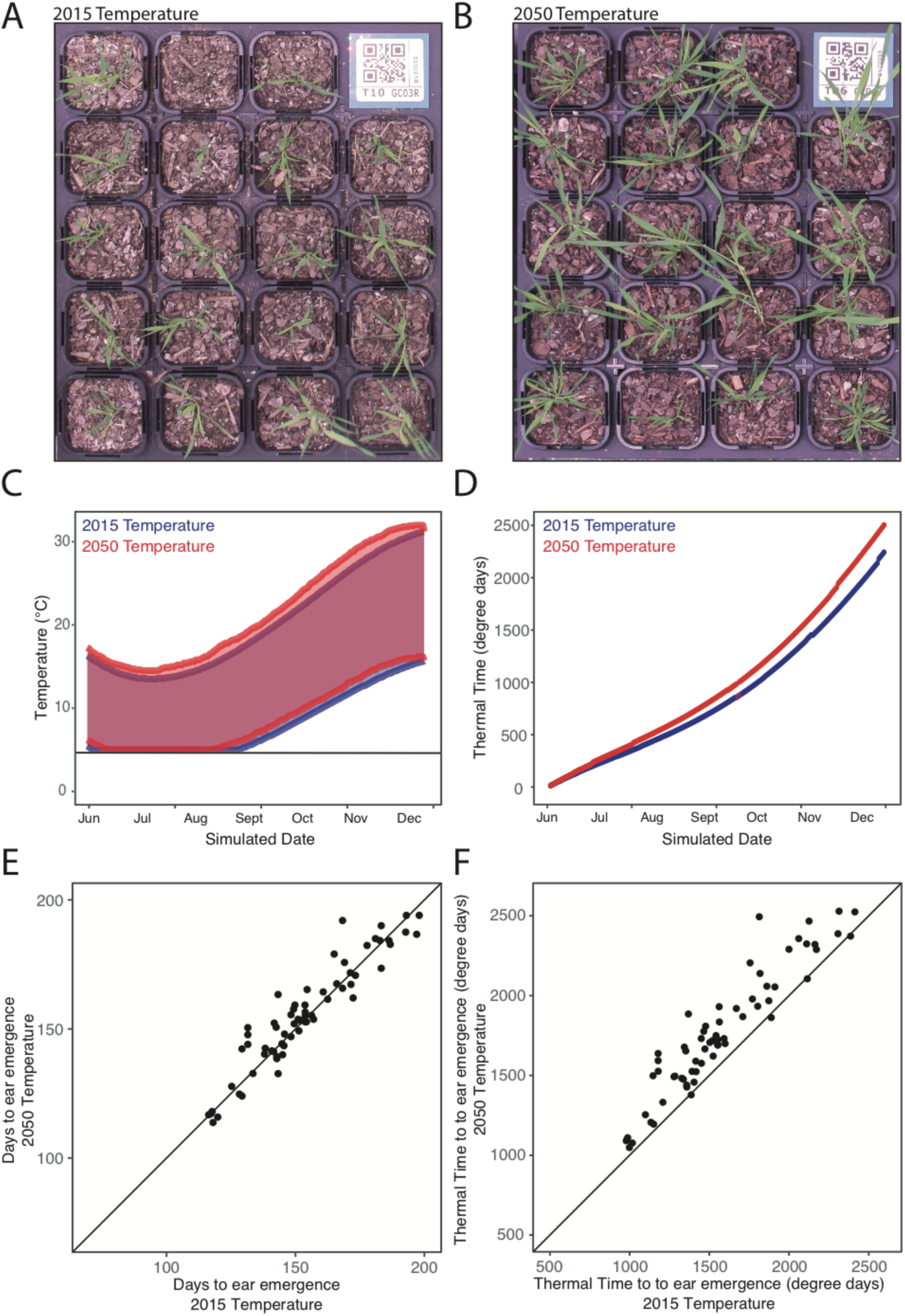
Climate chambers were used to compare the response of agronomic traits to small change in the climate for a Winter sowing in the Wagga Wagga region, south -eastern Australia. The GWAS HapMap set were grown in A) 2015 temperature climate and B) a 2050 temperature climate. Photos show representative plants after 16 weeks of growth. Climate chambers were programmed to have C) diurnal and seasonal changes in temperature resulting in different rates of accumulation of thermal time D) in the 2015 and 2050 climates. Timing of ear emergence was compared between chambers for both E) days to ear emergence and F) the accumulation of thermal time to ear emergence.

As expected, the accessions developed quicker and grew larger in the 2050 temperature profile (Fig 2A, B) as is consistent with a quicker accumulation of thermal time (Fig 2D). Early vigour parameters and energy use efficiency traits were measured when the majority of plants were at a four-leaf stage. Growth stages, tiller number and ear emergence date were monitored twice a week (Supp. S12, S13, S14). The experiment was ceased after 200 days of growth, at which time there were 5 and 7 lines that did not flower in the 2015 condition and 2050 conditions, respectively. The remaining lines reached ear emergence at a similar number of days in both the present and future conditions (Fig 2E). However, when converted to thermal time those lines in the 2015 temperature condition required less thermal time than those in the 2050 temperature condition (Fig 2D, F). This indicates that there is generally more dependence on photoperiod in this population than on thermal time to trigger the transition to flowering. There was variation between genotypes in the plasticity in their response to the two conditions (Fig 2E & F), indicating that it would be worthwhile mapping the genotype by environment interaction.

### Determining the genetic architecture of early growth, ear emergence and energy use efficiency traits in response to environment

GWAS was performed on raw and derived traits as described in the methods (Fig 3; Supp. S15 & S16). For ear emergence, eight significant QTLs were identified. EarEmerg_QTL3.1 explains 62% of the phenotypic variation in thermal time to ear emergence in the 2015 temperature condition while two QTLs, EarEmerg_QTL3.1 and EarEmerg_QTL5.3, explain 56% and 10% of the phenotypic variance in thermal time to ear emergence in the 2050 temperature condition, respectively. No QTLs were found to be significant in both conditions but EarEmerg_QTL5.3 was significant in the 2050 temperature condition and was just under the significant threshold in the 2015 temperature condition (Fig 4A, Supp. S17). Within the 100kb region of this SNP there are 15 genes of which several could be relevant to the regulation of flowering including a YABBY transcription factor (Bradi5g16910), a no apical meristem (NAM) protein (Bradi5g16917) and an expressed gene containing a RNA recognition motif (Bradi5g16930). Interestingly, there were two QTLs that were significant for thermal time to ear emergence, EarEmerg_QTL3.1 and EarEmerg_QTL4.2, but not for days to ear emergence. There were six QTLs identified for the genotype by environment interaction, explaining in part, the variation among lines in response to future climate.

**Figure 3.**
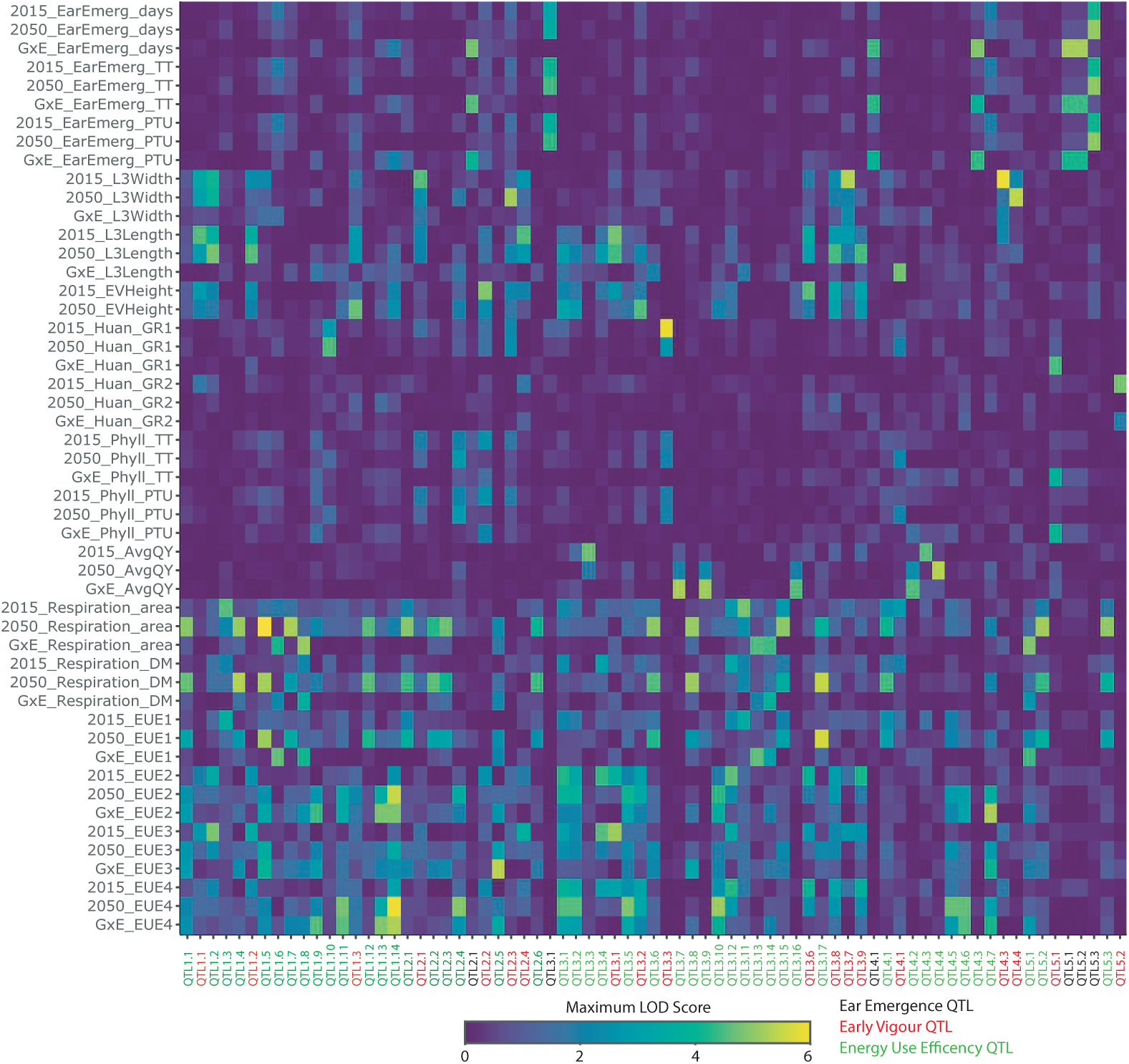
QTL were identified for a range of agronomic traits phenotyped in the 2015 temperature and 2050 temperature climates and the GxE interaction. A total of 73 significant QTL were identified by GWAS. There was little overlap between QTL for different traits but 2 robust QTL were identified in both environments while 16 QTL were identified for a genotype by environment (GxE) interaction. GxE - genotype by environment interaction; EarEmerg - ear emergence; TT - thermal time; PTU - photothermal units; L3Width - leaf 3 width; L3Length - leaf 3 length; GR - growth rate; GR - growth rate; EV - early vigour; phyll - phyllacron interval; AvgQY - average quantum yield; DM - dry mass; EUE - energy use efficiency

**Figure 4.**
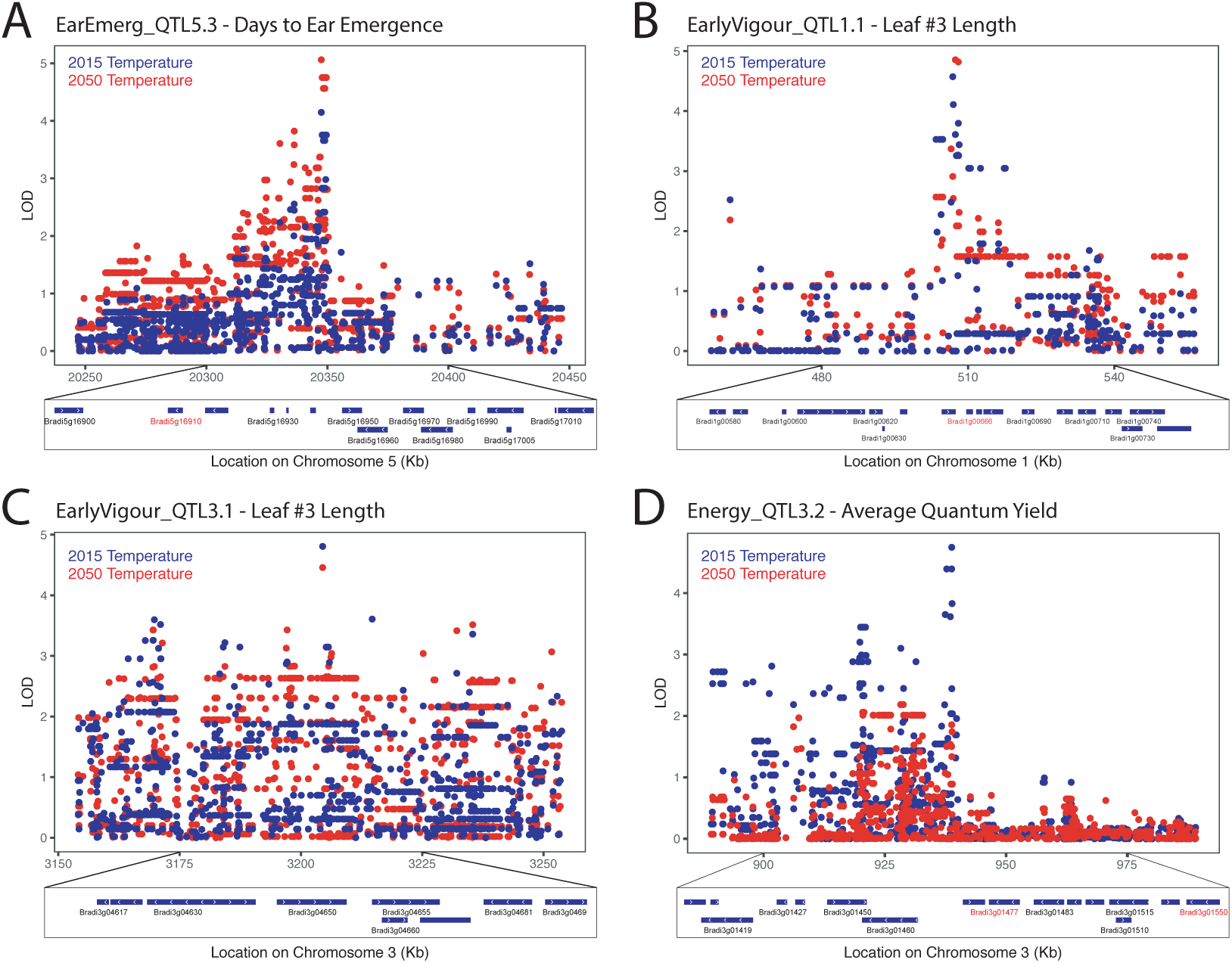
Putative candidate genes were identified under QTL of key interest. The ear emergence QTL, EarEmerg_QTL5.3, was significant for days to ear emergence in the 2050 temperature condition and only just under the significance threshold for the 2015 condition. Likely candidate genes include a YABBY transcription factor Bradi5g16910. B) The early vigour QTL, EarlyVigour_QTL1.1 for leaf #3 length was found to be significant in both conditions. This region contains an ethylene sensitive transcription factor, Bradi1g00666. C) The early vigour QTL, EarlyVigour_QTL3.1 was also identified for leaf #3 length in both environments. D) A strong QTL was identified for photosynthetic efficiency, Energy_QTL3.2, which was significant only in the 2015 temperature condition. Likely candidate genes include a heat shock protein, Bradi3g01477, and a Low Photosystems II Accumulation 3 (LPA3) protein, Bradi3g01550.

For early vigour, 22 significant QTLs were identified for 5 traits across the two climate conditions (Supp. S15). Two QTLs were identified in both conditions, EarlyVigour_QTL1.1 and EarlyVigour_QTL3.1, and both of these were for leaf #3 length. The 100kb region surrounding these QTLs contained 19 and 13 genes, respectively (Fig 4B and C). There was a highly significant QTL on chromosome 3 for growth rate 1, EarlyVigour_QTL3.3, a measure of the rate of development of the seedling at the two leaf stage, but only in the 2015 temperature condition. The 100kb region surrounding this QTL contained 13 genes (Supp. S18). A total of 6 QTLs were identified for the genotype x environment interaction across the two conditions for early vigour traits.

For the energy use efficiency traits, a total of 47 QTLs were identified across the two conditions for the three measured traits and four derived traits (Supp. S15). Of these QTL, none were found in both environments. However, a strong QTL, Energy_QTL3.3, was identified for average quantum yield, a measure of photosynthetic efficiency, in the 2015 temperature environment. The 100kb region around this QTL contained 24 genes including a low PSII accumulation 3 chloroplastic protein (Bradi3g01550), a Heat Shock Protein (Bradi3g01477) and several transcription factors (Fig4D, Supp. S19).

## Discussion

Thanks to the international *Brachypodium* community, in addition to our own collections, here we were able to provide the most comprehensive survey of *Brachypodium* species complex diversity to date. With 1897 accessions across the globe this is a greater than 10-fold increase from previous studies (25,32).

Since being described as three separate species in 2012 (22), species identification in the *Brachypodium* species complex has been achieved by morphology, PCR of a select set of markers or DNA barcoding (e.g. 41,42). Here we present a unique systematic method of determining the species of an accession using low coverage genotyping by sequencing and bioinformatics, providing a high throughput and low cost alternative for species identification. We found that the majority of our accessions were in fact *B. hybridum* (56%) including the vast majority of accessions in Australia and North America (Fig 1A). The wide dispersion of this species may be due to the benefit of the multiple genomes resulting from polyploidisation (47). There were relatively few *B. stacei* (3%), which were limited to the Mediterranean region (Fig 1A).

Within *B. distachyon* itself we found significant population structure including high level subspecies splits, with 6.5% divergence between subspecies, which is greater than that found between *indica* and *japonica* rice at 1.4% divergence (48). While many previous studies have focused on individual regions (32,33), the collection of 490 high quality *B. distachyon* accessions combined with 81,400 high quality SNPs presented here has allowed us to further distinguish subgroups with the *B. distachyon* subspecies, with and eastern and western European group in each subspecies. A number of geographically diverse highly related genotypic lineages were also identified which showed within lineage divergence of between 0.1- to 0.4% of SNPs. The geographic spread of these lineages highlights the inbreeding nature and high dispersal ability of *B. distachyon*.

The hierarchical levels of genetic variation within the *Brachypodium* species complex can be attributed to allopolyploidisation and subspeciation, possibly during the most recent ice age; east/west isolation by distance in Europe; and the high levels of self-fertilisation in the species (21). These levels of population structure have been seen in Arabidopsis (9), and other highly selfing plant species such as barley (49), but are more extreme in *Brachypodium*. In rice, either the *indica* sub-species (50) or *japonica* sub-species (51), were separately used for GWAS. Similarly, to deal with the population structure in this study, the HapMap set was limited to the A subspecies of *B. distachyon* with remaining relatedness included in the GWAS analysis using mixed models (QTLrel; 47).

The lack of recombinant genetic diversity with subspecies and populations of *B. distachyon* also limits the power of GWAS analysis. The HapMap set contains a large amount of genomic diversity (>1% of bases are variable) but the sample size is low and the extent of linkage disequilibrium is high, limiting mapping resolution. However, the patterns are similar to rice where GWAS is very effective as sample size increases (10). The construction of a Nested Association Mapping (NAM) population for *B. distachyon* would be advantageous to break-up the population and familial lineages and to increase the frequency of minor alleles. This has been a successful approach in other species such as maize and wheat (53,54).

In field conditions, determining the relationship between various physiological traits and their impact on yield is difficult due to in-season environmental variability and the presence of a range of abiotic and biotic stresses. However, experiments in growth chambers often have little relevance to field conditions due to the unrealistic and static nature of the conditions. The use of climate chambers provides a realistic range of temperatures, which results in more translatable results to the field (4,55), while removing the variability caused by weather and stresses. The use of climate chambers also allows the impact of small changes in climate to be observed and the dissection of which components of the climate have the largest influence on a trait of interest. In this study, we examined the effect of an increase in temperature in line with climate change model predictions for 2050 in south eastern Australia. Unexpectedly, there was generally a short delay of flowering time in the 2050 temperature condition with variation in the extent of delay in different genotypes while there was little dependence of flowering on the accumulation of thermal time. This indicates that there may be some vernalisation requirements in *B. distachyon* that are not being met in the 2050 temperature condition. The lack of vernalisation is also evident in the fact that seven lines had not flowered by the end of the 2050 temperature condition while 5 lines did not flower in the 2015 temperature condition. While this GWAS analysis did not identify known flowering time loci that regulate vernalisation-induced flowering such as VRN1, VRN2 and FT (37,56), the QTL may represent more subtle vernalisation processes that would be important for facultative varieties. Perhaps largely to the difference in growth conditions, the QTL in this study did not overlap with those found in a previous GWAS of flowering time (25). Candidate genes identified for flowering time here included several transcription factors, including a YABBY transcription factor whose closest ortholog in rice, Os04g45330, is most highly expressed in the shoot apical meristem and developing inflorescence (Rice Gene Expression Atlas) and whole closest ortholog in Arabidopsis, At2g45190, is involved in regulation of the floral morphology (57) under EarEmerg_QTL5.3. This QTL was significant in the 2050 temperature conditions and was only just below the significance threshold in the 2015 temperature condition (Fig 4A).

Early vigour is an important trait in many parts of Australia, and the world, where there is competition from weeds and a shorter season. Despite the highest correlating non-destructive measure of early above ground biomass being seedling height, the most robust QTLs across environments were actually identified by leaf #3 length. Two QTLs identified for leaf #3 length were identified in both environments, indicates they could potentially be useful for breeding for early vigour in multiple environment types. One of these, EarlyVigour_QTL1.1 is located in an area of synteny to other areas where early vigour QTLs have been identified at the end of chromosome 3 in rice (16,58,59) and Chromosome 4 in wheat (60). Within EarlyVigour_QTL1.1 there is a candidate gene, Bradi1g00666, that is described as an ethylene-responsive transcription factor. The main candidate gene in the QTL on Chromosome 3 in rice was also an ethylene responsive gene (58). The EarlyVigour_QTL3.1 for leaf #3 length was also found to be significant across both environments. There were no obvious candidate genes for this QTL but a number of signalling proteins that could be involved in molecular control of leaf size (Fig 4C, Supp. S18).

The balance of energy production and use in plants is highly linked to the conditions that the plant is grown under, however genetic variation controlling the energy efficiency of plants could be used to increase yield potentials. The quantum yield is an indicator of photosynthetic efficiency, the proportion of energy harvested through the light harvesting complexes that goes towards producing photosynthates (61). No QTL were identified in common across both environments, but there were 11 QTL that were identified for the GxE interaction. This may be due to the sensitivity of these energy processes to the subtle difference in environments or a result of being measured on different days to allow comparison of plants at the same developmental stage. A strong QTL was identified for quantum yield, a measure of the efficiency of photosystem II, in the 2015 climate but interestingly not in the 2050 climate. Candidate genes under this QTL included a gene with 66% homology to the Low PSII Accumulation 3 (LPA3) gene in Arabidopsis, which has been shown to be important in Photosystems II assembly (62). Further studies into the importance of this QTL in different conditions, as well as the other photosynthesis and respiration QTL, would be worthwhile.

In conclusion, the *Brachypodium* species complex is heavily structured at the ploidy, subspecies, population, and family levels. This limits the ability to identify the genetic basis of adaptation as relatively few recombinant genotypes were obtained. Despite these limitations, this study indicates the potential to use *Brachypodium distachyon*, a model for Pooideae grass crops, to identify genetic variation in key pathways underlying agricultural traits through Genome Wide Association Studies. Further wild collections and/or the development of a Nested Association Mapping (NAM) populations could address the limitation of recombinant genotypes and result in very high power mapping population typical of 1000 genome projects. As it now stands, *Brachypodium* is a good model for both polyploidisation, with likely multiple events among small divergent genomes, and for invasion biology with multiple widespread genotypes identified across continents, regions and sites.

## Materials and Methods

### Genotyping by Sequencing and Species Identification

Genotyping by sequencing (GBS) was undertaken as described by Elshire and colleagues (28) using PstI enzyme and a library of homemade barcoded adaptors (see https://github.com/borevitzlab/brachy-genotyping; 27,28). Approximately 384 samples were multiplexed to run on a single lane in an Illumina HiSeq 2000 with a median number of 564 000 100bp read pairs per sample (https://github.com/borevitzlab/brachy-genotyping). Sequencing runs were undertaken by the Biomolecular Resource Facility (JCSMR, ANU).

Axe (63) was used to demultiplex sequencing lanes into libraries, allowing no mismatches. AdapterRemoval (64) was used to remove contaminants from reads, and merge overlapping read pairs. Reads were aligned using BWA MEM (65,66) to the Bd21-3 (*B. distachyon*) and ABR114 (*B. stacei*) reference genomes (Phytozome v.12.1), and to a *B. hybridum* pseudo-reference genome created by concatenating the *B. stacei* and *B. distachyon* reference genomes (Supp. S01). Variants were called using the multiallelic model of samtools mpileup (67) and bcftools call (68). Variants were filtered with bcftools filter, keeping only SNPs of reasonable mapping and variant qualities (>=10) and sequencing depth across samples (>=5 reads across all samples).

To determine the species of each of the accessions, we computed the proportion of each Chromosome in the *B. hybridum* pseudo-reference covered with at least 3 reads, excluding reads which mapped to multiple locations in the pseudo-reference, using mosdepth (69). The proportions of the *B. distachyon/B. stacei genomes* covered were normalised to be in [0, 1], and then used to assign samples into threshold groups: *B. stacei* (< 0.03), intermediate *B. stacei / B. hybridum* (< 0.28), *B. hybridium* (<0.34), intermediate *B. hybridum / B. distachyon* (0.94) and *B. distachyon* (>0.94); an additional group consisted of low coverage samples (<100000 reads in total). Samples from intermediate and low coverage groups were excluded, and only variants in the respective genomes were used to allocate the three species groups.

### Population Structure of *B. distachyon*

To determine the population structure of *B. distachyon* a pairwise Identity By State genetic distance was calculated to identify among 490 high quality samples a core diversity set of 72 distinct genotypes using 82,800 SNPs derived from GBS data and the SNPRelate package using a z-score of 3.5. Occasionally, when genotypes are closely related, noise between technical replicates of an accession will result in them being split across the related genotypes. Therefore, we keep replicate(s) from the genotype with the majority of replicates for that accession, breaking ties by keeping the replicate with the lowest missing data. In addition, 29 accessions whose geographic origin was suspect were also excluded.

To avoid bias from including up to 30 inbred accessions of the same genotype, a reduced set was input into STRUCTURE V.2.3.4 (34). A total of six replicates were run of population (K) 1-13 with a burnin setting of 10,000 sets, and 100,000 permutations per run (Fig 1B, Supp. S04). The optimal K was determined as K=3 by Evanno’s Delta K, processed via Structure Harvester and CLUMPP (70–72). Barplots and pie charts were generated via in-house developed R scripts available through github (https://github.com/borevitzlab/brachy-genotyping-notes).

For *B. distachyon* the pairwise distance between genotypes was also calculated in R and plotted as a dendrogram (Supp. S02). From this a set of 107 accessions were selected to represent the genotypic diversity of the species for whole genome sequencing to maximise SNP coverage across the genome.

### Whole Genome Sequencing

For Whole Genome Sequencing (WGS), sequencing libraries for individual samples were prepared from 6 ng genomic DNA with the Nextera DNA Library Prep kit (Illumina, San Diego, CA, USA). Libraries were enriched and barcoded with custom i5-, and i7-compatible oligos and Q5® High-Fidelity DNA Polymerase (NEB, Ipswich, MA, USA). Libraries were pooled and sequenced in one lane on a NextSeq 500 sequencer (Illumina, San Diego, CA, USA).

Trimit (73) was used to clean WGS reads of adaptors, and merge overlapping read pairs. BWA MEM was then used to align these reads against the Bd21-1 reference genome (version 314_v3.1; 16). Variants were called using freebayes (74) with default parameters. Variants were filtered such that only variants meeting the following criteria were kept: variant quality >20, minor allele frequency >= 2%. Heterozygous variant calls were changed to missing; due to the inbred nature of these accessions, heterozygous calls were almost certainly erroneous. https://github.com/borevitzlab/brachy-genotyping Linkage Disequilibrium (LD) was calculated across the *B. distachyon* genome using consecutive windows of 2000 SNPs from the whole genome data of the HapMap 74 set (http://github.com/borevitzlab/brachy-genotyping-notes).

### Plant Growth

Individual grain of each genotype were planted 2.5cm deep in square plastic pots (5cm width, 8cm deep) in a mix of 50:50 soil:washed river sand which had been steam pasteurised. Pots were then placed at 4oC in the dark for three days to stratify the seed before being moved to specially modified climate chambers (see 4). In brief, these chambers have been fitted with 7 LED light panels and are controlled to change the light intensity, light spectrum, air temperature and humidity every 5 minutes. Climatic conditions were modelled using SolarCalc software (75). The Wagga Wagga region was centred around -35S, 147E with an elevation of 147m. Plants were fertilised with Thrive (N:P:K 25:5:8.8 + trace elements, Yates) and watered with tap water as needed. Growth stages were recorded based on the Huan developmental stage (76) up until stem elongation and thereafter the Zadoks scale was used. Total leaf area was measured with a Li-1300 Area Meter (Li-COR). For dry weight, leaf tissue was dried in a paper envelope at 60oC for five days before weighing.

### Conversions of Phenotypic Data

Thermal time was calculated from the logged condition within each chamber with the following formula:

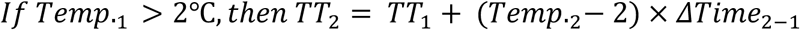

where *TT_i_* is accumulated thermal time at a particular timepoint *i* and *Temp_.i_* is the air temperature at a particular timepoint *i*.

Photothermal units (PTU) were calculated using the logged data from a PAR sensor in the middle of the chamber and the following formula:

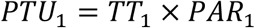

where TT is the accumulated thermal time at timepoint *i* and PAR is the measured photosynthetically active radiation at timepoint *i.*

Growth rates (GR) were calculated as:

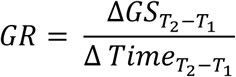

where GS is the Huan Growth Stage and T_1_ was approximately one leaf for the initial linear growth stage (GR1) T_1_ was approximately one leaf and three leaves and the faster growth stage (GR2) between three leaves and five leaves. The Phyllachron interval, the time taken to grow one leaf was calculated as:

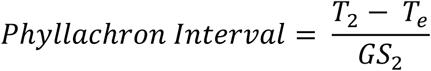

where T_2_ is the unit of time at approximately three leaf stage and T_e_ is the unit of time at seedling emergence for that particular plant. GS_2_ is the Huan Growth Stage at T_2_.

Final growth efficiency was calculated when plants reached ear emergence. The final growth efficiency 1 was calculated as:

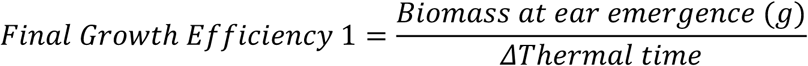

where accumulated thermal time is calculated from seedling emergence to ear emergence. The final growth efficiency 2 was calculated as:

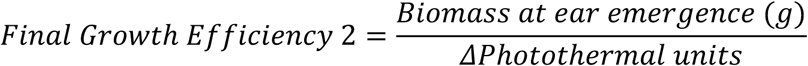

where accumulated photothermal units is calculated from seedling emergence to ear emergence.

### Energy Use Efficiency Traits

Energy use efficiency traits were measured on plants from the 2015-2050 Temperature experiment at a 4-5 leaf stage. Photosynthetic parameters were measured using a Trayscan system (PSI) incorporating Pulse Amplitude Modification (PAM) Chlorophyll Fluorescence measures of Quantum Efficiency (61). The parameters measured included photosynthetic efficiency, non-photochemical quenching and photo-inhibition. See Supp. S20 for Protocol.

Dark respiration rate was measured using the Q2 system (Astec Global) as in Scafaro et al (77). In brief, this system uses an oxygen sensitive fluorescent dye embedded in a cap to monitor the oxygen depletion with a tube containing the sample. A 3cm fragments in the centre of the last fully expanded leaf of each plant was used to measure dark respiration per unit area and per unit dry mass.

Several energy use efficiency formulas were calculated. These included a ratio of dark respiration to photosynthesis and measures of growth per unit dark respiration. These were as follows:

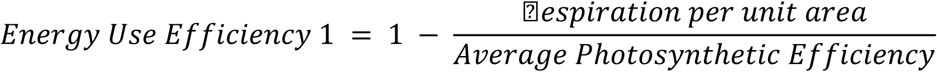

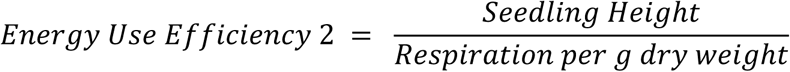

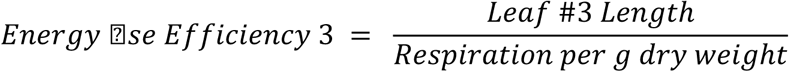

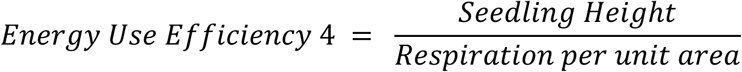

### Heritability

Narrow-sense heritability was calculated from the phenotype data using the nlme package in R.

### GWAS analysis

In preparation for GWAS, the genotype data was filtered to remove non-variant SNPs and redundant SNPs (i.e. SNPs whose genotypes are not different from adjacent SNPs but have more missing data points). Then SNPs with a minor allele frequency of < 3% were filtered out. As there was 18.5% missing data in the original data set, imputation was undertaken. First, if the observed genotypes of two adjacent SNPs were not different, then the missing genotype of one SNP was replaced by the observed genotype of the other SNP. Secondly, nearest neighborhood (NN) method was implemented to impute the remaining missing genotypes based on Huang et al (78) with some modifications. The nearest 50 SNPs from each side of the SNP under imputation were selected to estimate similarity between each pair of accessions, and then the missing genotype of an accession was replaced by the observed majority genotype of the closest 5 accessions. These parameters were determined by simulations to achieve an optimal imputation success rate, which was 97.95% for our data. Finally, SNPs with a minor allele frequency <5% were filtered.

Linear mixed-effect models were employed to identify genetic variants underlying phenotypes of interest

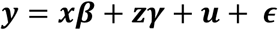

where ***y*** = (*y1, y2, ……., yn*)^’^ denotes phenotypic values, ***x*** = (*xij)nx(k+1)* represents intercept and *k* covariates (if any) with effects **β**, ***z*** is a vector of the coded genotypes at a scanning locus with effect γ, ***u*** = (*u1, u2, ……, un*)’ represents polygenic variation, and ***ϵ*** = (*ϵ1, ϵ2, ……, ϵn*) the residual effect. It was assumed that

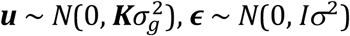
 and ***u*** was independent of ***ϵ***. The genetic relationship matrix ***K*** was estimated by identify-by-state (IBS) from genotypic data with markers on the Chromosome under scan being excluded to avoid proximal contamination (79,80). Estimation of ***K*** and genome scan were performed in R package QTLRel (52).

To determine a significance threshold, the permutation test was implemented on 1000 permutations of the phenotype data to estimate the genome-wide significance threshold at 0.05 for the trait of days to ear emergence. The significance threshold was determined to be a LOD (Logarithm of ODds) of 4.43583.

## Acknowledgements

*Brachypodium* Accession Contributors:ShuangShuang Liu, Kent Bradford PI, Smadar Ezrati PI, Hikmet Budak PI, Diana Lopez, Pilar Catalan PI, David Garvin PI, John Vogel PI, Sean Gordon, Sam Hazen PI, Luis Mur PI We would like to acknowledge the technical assistance of Suyan Yee and Allison Heussler.

## Supplementary Materials

S01. Coverage of *B. stacei* and *B. distachyon* Genomes for Species ID

S02. Structure of *B. distachyon* collection as sequenced by GBS

S03. Dendrogram of reduced *B. distachyon* set used for STRUCTURE

S04. *B. distachyon* reduced set STRUCTURE plots for k=2

S05. Linkage disequilibrium across the *B. distachyon* genome

S06. Graphical representation of climatic conditions from the flowering time experiment

S07. Phenotype data from flowering time experiment

S08. Flowering time experiment across 266 *B. distachyon* accessions

S09. Phenotypic correlation of traits from the early vigour validation experiment

S10. List of HapMap 74 genotypes

S11. Assessment of climate change models for South East Australia

S12. Phenotype data from 2015 temperature and 2050 temperature climate experiment

S13. Heritability of traits of interest from 2015 temperature and 2050 temperature climate experiment

S14. Histograms of all traits in 2015 temperature and 2050 temperature climate experiment

S15. List of QTL identified for traits of interest in 2015 temperature and 2050 temperature climate experiment

S16. Manhatten Plots for all traits in 2015 temperature and 2050 temperature climate experiment

S17. List of candidate genes for strong ear emergence QTL

S18. List of candidate genes for strong early vigour QTL

S19. List of candidate genes for strong energy QTL

S20. Protocol for measuring phenotypic parameters using a PAM

## References

Wheeler T, von Braun J. Climate Change Impacts on Global Food Security. Science (80-) [Internet]. 2013 Aug 2;341(6145):508 LP-513. Available from: http://science.sciencemag.org/content/341/6145/508.abstract

Ray DK, Mueller ND, West PC, Foley JA. Yield Trends Are Insufficient to Double Global Crop Production by 2050. PLoS One. 2013;8(6):e66428.

Rivers J, Warthmann N, Pogson BJ, Borevitz JO. Genomic breeding for food, environment and livelihoods. Food Secur [Internet]. 2015 Apr;7(2):375–82. Available from: https://doi.org/10.1007/s12571-015-0431-3

Brown TB, Cheng R, Sirault XRR, Rungrat T, Murray KD, Trtilek M, et al. TraitCapture: genomic and environment modelling of plant phenomic data. Curr Opin Plant Biol [Internet]. 2014;18(Supplement C):73–9. Available from: http://www.sciencedirect.com/science/article/pii/S1369526614000181

Condon AG, Richards RA, Rebetzke GJ, Farquhar GD. Breeding for high water-use efficiency. J Exp Bot. 2004;55:2447–60.

Wilson PB, Rebetzke GJ, Condon AG. Of growing importance:combining greater early vigour and transpiration efficiency for wheat in variable rainfed environments. Funct Plant Biol. 2015;42:1107–15.

Wilson D, Jones J. Effect of selection for dark respiration rate of mature leaves on crop yields of Lolium perenne cv. S23. Ann Bot [Internet]. 1982;49:313–20. Available from: http://aob.oxfordjournals.org/content/49/3/313.short

Winzeler M, Mccullough DE, Hunt LA. Genotypic Differences in Dark Respiration of Mature Leaves. Can J Plant Sci. 1988;68:669–75.

Atwell S, Huang YS, Vilhjalmsson BJ, Willems G, Horton M, Li Y, et al. Genome-wide association study of 107 phenotypes in Arabidopsis thaliana inbred lines. Nature. 2010;465(7298):627–31.

Huang X, Han B. Huang2014 - Crop GWAS review. Annu Rev Plant Biol. 2014;65:531–51.

Sukumaran S, Dreisigacker S, Lopes M, Chavez P, Reynolds MP. Genome-wide association study for grain yield and related traits in an elite spring wheat population grown in temperate irrigated environments. Theor Appl Genet. 2014;128(2):353–63.

Garvin DF, Gu YQ, Hasterok R, Hazen SP, Jenkins G, Mockler TC, et al. Development of genetic and genomic research resources for Brachypodium distachyon, a new model system for grass crop research. Vol. 48, Crop Science. 2008. p. S69–84.

Brutnell TP, Bennetzen JL, Vogel JP. Brachypodium distachyon and Setaria viridis: Model Genetic Systems for the Grasses. Annu Rev Plant Biol. 2015;66:465–85.

Draper J, Mur LAJ, Jenkins G, Ghosh-Biswas GC, Bablak P, Hasterok R, et al. Brachypodium distachyon. A New Model System for Functional Genomics in Grasses. Plant Physiol [Internet]. 2001;127(4):1539–55. Available from: http://www.plantphysiol.org/content/127/4/1539.abstract

Mur LAJ, Allainguillaume J, Catalan P, Hasterok R, Jenkins G, Lesniewska K, et al. Exploiting the Brachypodium Tool Box in cereal and grass research. New Phytol. 2011;191(2):347.

The International Brachypodium Initiative. Genome sequencing and analysis of the model grass Brachypodium distachyon. Nature [Internet]. 2009;70(1–2):47–61. Available from: http://link.springer.com/10.1007/s11103-009-9456-3

The International Wheat Genome Sequencing Consortium [Internet]. 2017. Available from: www.wheatgenome.org

The International Barley Genome Sequencing Consortium. A physical, genetic and functional sequence assembly of the barley genome. Nature. 2012;491:711.

Wicker T, Mayer KFX, Gundlach H, Martis M, Steuernagel B, Scholz U, et al. Frequent Gene Movement and Pseudogene Evolution Is Common to the Large and Complex Genomes of Wheat, Barley, and Their Relatives. Plant Cell [Internet]. 2011;23(5):1706–18. Available from: http://www.plantcell.org/lookup/doi/10.1105/tpc.111.086629

Vogel JP, Tuna M, Budak H, Huo N, Gu YQ, Steinwand MA. Development of SSR markers and analysis of diversity in Turkish populations of Brachypodium distachyon. BMC Plant Biol. 2009;9:88.

Wilson PB, Streich JC, Borevitz JO. Genomic Diversity and Climate Adaptation in Brachypodium. In: Vogel J, editor. Genetics and Genomics of Brachypodium. Switzerland: Springer International Publishing; 2015. p. 107–28.

Catalan P, Muller J, Hasterok R, Jenkins G, Mur LAJ, Langdon T, et al. Evolution and taxonomic split of the model grass Brachypodium distachyon. Ann Bot. 2012;109(2):385–405.

Hasterok R, Draper J, Jenkins G. Laying the cytotaxonomic foundations of a new model grass, Brachypodium distachyon (L.) beauv. Chromosom Res. 2004;12(4):397–403.

Idziak D, Hazuka I, Poliwczak B, Wiszynska A, Wolny E, Hasterok R. Insight into the karyotype evolution of Brachypodium species using comparative chromosome barcoding. PLoS One. 2014;9(3).

Tyler L, Lee SJ, Young ND, DeIulio GA, Benavente E, Reagon M, et al. Population Structure in the Model Grass Is Highly Correlated with Flowering Differences across Broad Geographic Areas. Plant Genome [Internet]. 2016;9(2):0. Available from: https://dl.sciencesocieties.org/publications/tpg/abstracts/9/2/plantgenome2015.08.0074

Gordon SP, Contreras-Moreira B, Woods DP …, Vogel JP. Extensive gene content variation in the Brachypodium distachyon pan-genome correlates with population structure. Nat Commun. 2017;2184.

Catalan P, Lopez-Alvarez D, Bellosta C, Villar L. Updated taxonomic descriptions, iconography, and habitat preferences of Brachypodium distachyon, B. stacei, and B. hybridum (Poaceae). An del Jard Bot Madrid. 2016;73(1):e028.

Elshire RJ, Glaubitz JC, Sun Q, Poland JA, Kawamoto K, Buckler ES, et al. A Robust, Simple Genotyping-by-Sequencing (GBS) Approach for High Diversity Species. PLoS One [Internet]. 2011;6(5):e19379. Available from: http://dx.doi.org/10.1371/journal.pone.0019379

Morris GP, Grabowski PP, Borevitz JO. Genomic diversity in switchgrass (Panicum virgatum): From the continental scale to a dune landscape. Mol Ecol. 2011;20(23):4938–52.

Nicotra AB, Chong C, Bragg JG, Ong CR, Aitken NC, Chuah A, et al. Population and phylogenomic decomposition via genotyping-by-sequencing in Australian Pelargonium. Mol Ecol. 2016;25(9):2000–14.

Zheng X, Levine D, Shen J, Gogarten SM, Laurie C, Weir BS. A high-performance computing toolset for relatedness and principal component analysis of SNP data. Bioinformatics. 2012;28(24):3326–8.

Filiz E, Ozdemir BS, Budak F, Vogel JP, Tuna M, Budak H. Molecular, morphological, and cytological analysis of diverse Brachypodium distachyon inbred lines. Genome. 2009;52(10):876–90.

Marques I, Shiposha V, López-Alvarez D, Manzaneda AJ, Hernandez P, Olonova M, et al. Environmental isolation explains Iberian genetic diversity in the highly homozygous model grass Brachypodium distachyon. BMC Evol Biol. 2017;17(1).

Pritchard JK, Stephens M, Donnelly P. Inference of population structure using multilocus genotype data. Genetics. 2000;

Brachi B, Faure N, Horton M, Flahauw E, Vazquez A, Nordborg M, et al. Linkage and association mapping of Arabidopsis thaliana flowering time in nature. PLoS Genet. 2010;

Maurer A, Draba V, Jiang Y, Schnaithmann F, Sharma R, Schumann E, et al. Modelling the genetic architecture of flowering time control in barley through nested association mapping. BMC Genomics [Internet]. 2015;16(1):290. Available from: http://www.biomedcentral.com/1471-2164/16/290

Bettgenhaeuser J, Corke FMK, Opanowicz M, Green P, Hernández-Pinzón I, Doonan JH, et al. Natural Variation in Brachypodium Links Vernalization and Flowering Time Loci as Major Flowering Determinants. Plant Physiol [Internet]. 2017;173(1):256–68. Available from: http://www.plantphysiol.org/lookup/doi/10.1104/pp.16.00813

Higgins JA, Bailey PC, Laurie DA. Comparative Genomics of Flowering Time Pathways Using Brachypodium distachyon as a Model for the Temperate Grasses. PLoS One. 2010;5(4).

Ream TS, Woods DP, Schwartz CJ, Sanabria CP, Mahoy J a, Walters EM, et al. Interaction of photoperiod and vernalization determines flowering time of Brachypodium distachyon. Plant Physiol [Internet]. 2014;164(2):694–709. Available from: http://www.pubmedcentral.nih.gov/articlerender.fcgi?artid=3912099&tool=pmcentrez&rendertype=abstract

Woods DP, Bednarek R, Bouché F, Gordon SP, Vogel JP, Garvin DF, et al. Genetic Architecture of Flowering-Time Variation in Brachypodium distachyon. Plant Physiol [Internet]. 2017;173(1):269–79. Available from: http://www.ncbi.nlm.nih.gov/pubmed/27742753%0A http://www.plantphysiol.org/lookup/doi/10.1104/pp.16.01178

Rebetzke GJ, Richards RA. Genetic improvement of early vigour in wheat. Aust J Agric Res. 1999;50(3):291–301.

Wilson PB, Rebetzke GJ, Condon AG., Field Crops Research Pyramiding greater early vigour and integrated transpiration efficiency in bread wheat ; trade-offs and benefits. F Crop Res [Internet]. Elsevier B.V.; 2015;183:102–10. Available from: http://dx.doi.org/10.1016/j.fcr.2015.07.002

Brachi B, Morris GP, Borevitz JO. Genome-wide association studies in plants: The missing heritability is in the field. Vol. 12, Genome Biology. 2011.

Climate Change in Australia website [Internet]. [cited 2015 Sep 1]. Available from: https://www.climatechangeinaustralia.gov.au/en/climate-projections/climate-futures-tool/introduction-climate-futures/

Giraldo P, Rodríguez-Quijano M, Vázquez JF, Carrillo JM, Benavente E. Validation of microsatellite markers for cytotype discrimination in the model grass Brachypodium distachyon. Genome. 2012;55(7):523–7.

López-Alvarez D, López-Herranz ML, Betekhtin A, Catalán P. A DNA barcoding method to discriminate between the model plant Brachypodium distachyon and its close relatives B. stacei and B. hybridum (Poaceae). PLoS One [Internet]. 2012;7(12):e51058. Available from: http://www.pubmedcentral.nih.gov/articlerender.fcgi?artid=3519806&tool=pmcentrez&rendertype=abstract

Te Beest M, Le Roux JJ, Richardson DM, Brysting AK, Suda J, Kubešová M, et al. The more the better? The role of polyploidy in facilitating plant invasions. Vol. 109, Annals of Botany. 2012. p. 19–45.

Ma J, Bennetzen JL. Rapid recent growth and divergence of rice nuclear genomes. Proc Natl Acad Sci [Internet]. 2004;101(34):12404–10. Available from: http://www.pnas.org/cgi/doi/10.1073/pnas.0403715101

Wang M, Jiang N, Jia T, Leach L, Cockram J, Waugh R, et al. Genome-wide association mapping of agronomic and morphologic traits in highly structured populations of barley cultivars. Theor Appl Genet. 2012;124(2):233–46.

Huang X, Zhao Y, Wei X, Li C, Wang A, Zhao Q, et al. Genome-wide association study of flowering time and grain yield traits in a worldwide collection of rice germplasm. Nat Genet [Internet]. 2012;44(1):32–9. Available from: http://dx.doi.org/10.1038/ng.1018

Yano K, Yamamoto E, Aya K, Takeuchi H, Lo PC, Hu L, et al. Genome-wide association study using whole-genome sequencing rapidly identifies new genes influencing agronomic traits in rice. Nat Genet [Internet]. 2016 Jun 20;48(8):927–34. Available from: http://www.nature.com/doifinder/10.1038/ng.3596

Cheng R, Abney M, Palmer AA, Skol AD. QTLRel: An R Package for Genome- wide Association Studies in which Relatedness is a Concern. BMC Genet. 2011;12.

Bajgain P, Rouse MN, Tsilo TJ, Macharia GK, Bhavani S, Jin Y, et al. Nested association mapping of stem rust resistance in wheat using genotyping by sequencing. PLoS One. 2016;11(5).

Tian F, Bradbury PJ, Brown PJ, Hung H, Sun Q, Flint-Garcia S, et al. Genome-wide association study of leaf architecture in the maize nested association mapping population. Nat Genet [Internet]. 2011;43(2):159–62. Available from: http://dx.doi.org/10.1038/ng.746

Poorter H, Fiorani F, Pieruschka R, Wojciechowski T, van der Putten WH, Kleyer M, et al. Pampered inside, pestered outside? Differences and similarities between plants growing in controlled conditions and in the field. Vol. 212, New Phytologist. 2016. p. 838–55.

Woods DP, Ream TS, Amasino RM., Memory of the vernalized state in plants including the model grass Brachypodium distachyon. Front Plant Sci. 2014;5(March):99.

Boter M, Golz JF, Giménez-Ibañez S, Fernandez-Barbero G, Franco-Zorrilla JM, Solano R. FILAMENTOUS FLOWER Is a Direct Target of JAZ3 and Modulates Responses to Jasmonate. Plant Cell [Internet]. 2015;27(11):3160–74. Available from: http://www.plantcell.org/lookup/doi/10.1105/tpc.15.00220

Singh UM, Yadav S, Dixit S, Ramayya PJ, Devi MN, Raman KA, et al. QTL Hotspots for Early Vigor and Related Traits under Dry Direct-Seeded System in Rice (Oryza sativa L.). Front Plant Sci [Internet]. 2017;8:286. Available from: https://www.frontiersin.org/article/10.3389/fpls.2017.00286

Lu XL, Niu AL, Cai HY, Zhao Y, Liu JW, Zhu YG, et al. Genetic dissection of seedling and early vigor in a recombinant inbred line population of rice. Plant Sci. 2007;172(2):212–20.

Rebetzke GJ, Appels R, Morrison AD, Richards RA, McDonald G, Ellis MH, et al. Quantitative trait loci on chromosome 4B for coleoptile length and early vigour in wheat (Triticum aestivum L.). Aust J Agric Res. 2001;52(11–12):1221–34.

Rungrat T, Awlia M, Brown T, Cheng R, Sirault X, Fajkus J, et al. Using Phenomic Analysis of Photosynthetic Function for Abiotic Stress Response Gene Discovery Using Phenomic Analysis of Photosynthetic Function for Abiotic Stress Response Gene Discovery. Arab B. 2016;e0185.

Lu Y. Identification and Roles of Photosystem II Assembly, Stability, and Repair Factors in Arabidopsis. Front Plant Sci [Internet]. 2016;7. Available from: http://journal.frontiersin.org/Article/10.3389/fpls.2016.00168/abstract

Murray KD, Borevitz JO. Axe: rapid, competitive sequence read demultiplexing using a trie. BioRxiv. 2017;

Schubert M, Lindgreen S, Orlando L. AdapterRemoval v2: Rapid adapter trimming, identification, and read merging. BMC Res Notes. 2016;9(1).

Li H. Aligning sequence reads, clone sequences and assembly contigs with BWA-MEM. arXiv. 2013;

Li H, Durbin R. Fast and accurate short read alignment with Burrows-Wheeler transform. Bioinformatics. 2009;25(14):1754–60.

Li H. A statistical framework for SNP calling, mutation discovery, association mapping and population genetical parameter estimation from sequencing data. Bioinformatics. 2011;27(21):2987–93.

Danecek P, Schiffels S, Durbin R. Multiallelic calling model in bcftools (-m) [Internet]. 2016. Available from: http://samtools.github.io/bcftools/call-m.pdf

Pedersen BS, Quinlan AR. Mosdepth: quick coverage calculation for genomes and exomes. Bioinformatics [Internet]. 2017; Available from: http://academic.oup.com/bioinformatics/advance-article/doi/10.1093/bioinformatics/btx699/4583630

Evanno G, Regnaut S, Goudet J. Detecting the number of clusters of individuals using the software STRUCTURE: A simulation study. Mol Ecol. 2005;14(8):2611–20.

Jakobsson M, Rosenberg NA. CLUMPP: A cluster matching and permutation program for dealing with label switching and multimodality in analysis of population structure. Bioinformatics. 2007;23(14):1801–6.

Earl DA, vonHoldt BM. STRUCTURE HARVESTER: A website and program for visualizing STRUCTURE output and implementing the Evanno method. Conserv Genet Resour. 2012;4(2):359–61.

Murray KD, Borevitz JO. libqcpp: A C++14 sequence quality control library. J Open Source Softw. 2017;2(13):232.

Garrison E, Marth G. Haplotype-based variant detection from short-read sequencing. 2012;

Spokas K, Forcella F. Estimating hourly incoming solar radiation from limited meteorological data. Weed Sci [Internet]. 2006;541:182–9. Available from: https://www.cambridge.org/core/product/identifier/S0043174500007670/type/journal_article

Haun JR. Visual Quantification of Wheat Development. Agron J. 1973;65:116–9.

Scafaro AP, Negrini ACA, O’Leary B, Rashid FAA, Hayes L, Fan Y, et al. The combination of gas-phase fluorophore technology and automation to enable high-throughput analysis of plant respiration. Plant Methods. 2017;13(1).

Huang X, Wei X, Sang T, Zhao Q, Feng Q, Zhao Y, et al. Genome-wide association studies of 14 agronomic traits in rice landraces. Nat Genet [Internet]. 2010;42(11):961–7. Available from: http://dx.doi.org/10.1038/ng.695

Listgarten J, Lippert C, Kadie CM, Davidson RI, Eskin E, Heckerman D. Improved linear mixed models for genome-wide association studies. Vol. 9, Nature Methods. 2012. p. 525–6.

Cheng R, Parker CC, Abney M, Palmer AA. Practical Considerations Regarding the Use of Genotype and Pedigree Data to Model Relatedness in the Context of Genome-Wide Association Studies. G3 Genes|Genomes|Genetics [Internet]. 2013;3(10):1861–7. Available from: http://g3journal.org/lookup/doi/10.1534/g3.113.007948

